# Impact of Adipogenic Differentiation on the Nutrition and Sensory Profile of Cultivated Pork Fat Tissue

**DOI:** 10.64898/2026.09.16.749166

**Authors:** Emily T. Lew, Natsu Sugama, Sam S. Magrath, Scott C. Frost, David L. Kaplan

## Abstract

Adipose tissue is a principal source of texture, flavor, and aroma in meat products. Higher fat content corresponds to higher juiciness, tenderness and overall palatability of meat products, making it a key component of cultivated meat. Our previous study investigated the aromatic characteristics of cultivated pork fat tissue, finding comparable consumer panel hedonic scores between conventional livestock derived and cultivated pork fats. Further research was pursued to characterize if adipogenic differentiation of the cells is required to enhance the flavor of cultivated fat. While differentiation of cultivated cells results in tissues that more closely resemble the morphology of animal-derived tissue, the process substantially increases production timelines and material costs associated with cultivated meat production. As a result, we observed changes in organoleptic and nutritional properties of cultivated pork fat cells associated with differentiation of porcine dedifferentiated fat (PDFAT) cells. We assessed the effect of varying degrees of adipogenesis with soybean oil supplementation on fat content, composition, and volatile compound formation. Notably, we observed the concentrations of 2-pentylfuran, 2-heptanone, hexanal and (E)-2-octenal increased compared to the undifferentiated preadipocytes. These compounds, along with other ketones, aldehydes, alcohol and lactone molecules, contribute to fruity, herbal, and fatty aromas. Additionally, these results were achieved without requiring the maximum differentiation time frame of 12 days in culture. These findings emphasize the importance of adipocyte differentiation in cultivated fat production and highlight its potential as a strategy for optimizing both flavor and nutritional attributes of cultivated meat.

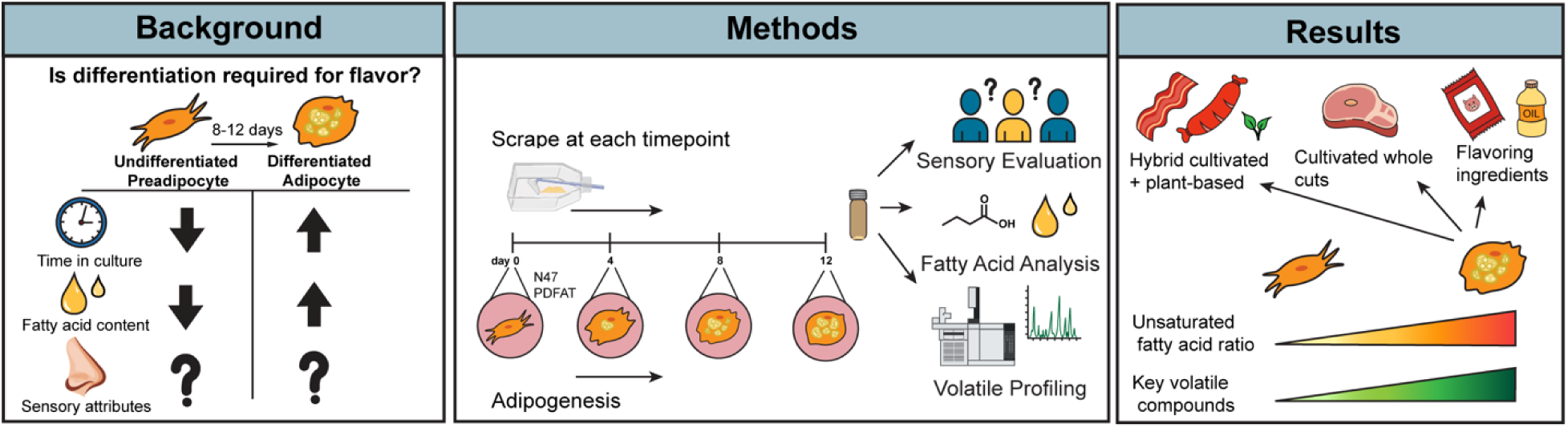

## Introduction

The projected demand for animal-sourced food products has led to rising concerns about the environmental impact of animal agriculture^1,2^. For example, global pork consumption has increased by 77% from 63.5 million tons in 1990 to 113 million tons in 2022^3^. This has increased demand for scarce resources such as land, water, and energy, and induced negative impacts on air, water and soil quality through greenhouse gas emissions and eutrophication^4–6^. In addition, the increasing demand for meat has raised concerns regarding public health and animal welfare^7,8^. Together, these challenges have fueled interest in sustainable food solutions such as plant-based meat and cultivated meat.

Plant-based meat options have expanded substantially, evolving beyond traditional staples such as tofu, tempeh, and seitan to include more advanced analogues intended to more closely replicate burgers, sausages, nuggets and other widely consumed meat products^9^. These products are commonly based on plant protein isolates or concentrates, particularly from pea and soy^10^. Despite their widespread availability, many consumers are often not purchasing these products due to their suboptimal flavor^11,12^. Retail data indicates low repeat purchase rates for plant-based meat products, with approximately 63% of consumers making only a single purchase, while only 37% make a second purchase^13^. Consumers noted flavor as the most influential factor in their purchasing decisions^13^. Cultivated meat has demonstrated the potential to close the flavor gap between plant-based meat and livestock grown meat^14,15^.

Cultivated meat, produced through the in vitro culture of animal cells, offers the potential to alleviate the negative environmental and public health impacts of conventional livestock production while also addressing the flavor limitations of plant-based alternatives^16–18^. The incorporation of cultivated fat tissues into plant-based meats enables the development of hybrid meat products, in which sustainable, cost-effective plant ingredients are supplemented with the complex, species-specific flavors derived from animal fat cells^19^. Studies have also explored the integration of cultivated cells with plant-based scaffolds to mimic the textural and nutritional profile of conventional meat^20,21^. Common plant proteins such as textured soy protein have been studied with bovine muscle cells, generating a product that more closely mimics the structural and mechanical properties of conventional beef. The inclusion of cultivated muscle cells in these products also improved the meaty flavor and sensory attractiveness^22^. Similarly, hybrid approaches with cultivated fat can make cultivated meat more accessible by addressing the key challenge of tissue scale-up, while also improving the flavor quality and consumer acceptance of plant-based proteins.

Flavor development in cooked meat is driven by multiple complex reactions which generate a wealth of volatile organic compounds (VOCs) ^23–25^. The main reactions that contribute to cooked meat flavor are the Maillard reaction between amino compounds and reducing sugars, lipid oxidation, vitamin degradation, and lipid-Maillard interaction between lipid-oxidized products with the products of the Maillard reaction^26^. The Maillard reaction produces diverse volatile compounds such as sulfur containing molecules, furans, pyrazines, and lipid degradation generates many aliphatic aldehydes, ketones, alcohols, acids, esters, and other compounds responsible for species specific meat aromas^23,27^. While these reactions are well documented in conventional meat, production of these key aroma and flavor volatiles in cultivated meat has been scarcely researched. Recent studies have reported on the general aroma profiles of cultivated fat, muscle and fibroblasts^28–31^. These studies found that cultivated fat and muscle contain a similar profile of aldehydes and ketones as conventional meat, but secondary lipid oxidation products provide a somewhat different profile^28–31^. Studies have also reported differences in amino acid content between cultivated and conventional meat, which heavily impact umami and meaty flavor attributes^32,33^. Culture media has been cited as a method to tune cultivated meat flavor^31,34^, but little is known about how to modulate cultivated meat flavor through cell maturity and biological processes such as differentiation. A recent study found the differentiation of bovine muscle and adipose-derived mesenchymal stem cells increases Maillard reaction-derived bitter almond and roasted beef aromas^35^, however detailed investigations into the role of differentiation in flavor formation remain limited.

To address this gap, we present an investigation of adipogenesis as a function of flavor development in cultivated porcine fat. Focusing primarily on lipid oxidation and degradation products, we explored if differentiation is needed for proper volatile production that would mimic conventional livestock fat. Our findings offer insights on how cultivated fat aroma and nutritional composition can be modulated through adipogenesis and how this process can be used to enhance the sensory qualities of alternative meat.

## Results

### N47 PDFAT achieves higher differentiation than PDFAT1 parent population

To select a suitable porcine dedifferentiated fat (PDFAT) cell line for this study, the adipogenic capacity of PDFAT1 primary cells and N47 cells were compared. N47 is a clonal population derived from PDFAT1 primary cells isolated from the Yorkshire pig (*Sus domesticus*)^28^. At passage 12, N47 and PDFAT1 were differentiated for up to 12 days with 2 days of adipogenic induction media and 10 days of lipid accumulation or maintenance media. Beginning at day 4, N47 achieved a higher percent differentiation than its parent population. Neither N47 nor PDFAT1 showed significant increases in percent differentiation between day 8 and day 12 (Figure 1A,C). Beginning at day 8, N47 achieved greater lipid accumulation than PDFAT1, shown by higher normalized BODIPY intensity. N47 also demonstrated higher lipid accumulation than PDFAT1 through day 12 (Figure 1B). Both cell lines have multilocular phenotype (Figure 1D). N47 was selected to complete the chemical and sensory analysis.

**Figure 1.**
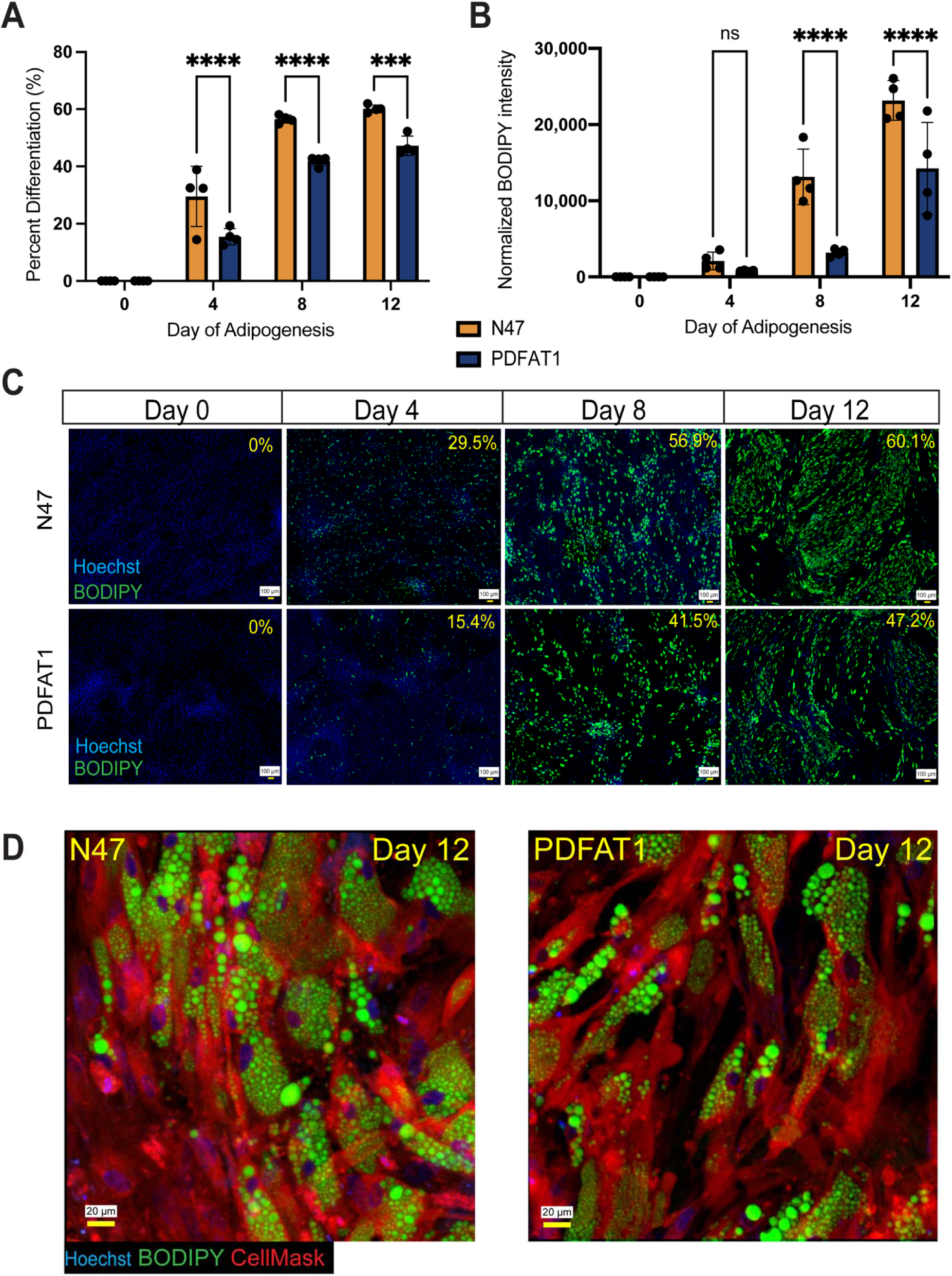
Comparison of differentiation capacity of N47 PDFAT clonal cell line and PDFAT1 primary PDFAT population. (A) differentiation percentage, (B) differentiation intensity and (C) morphology are shown at harvesting timepoints. Differentiation percent is shown in upper right corner. Scale bar denotes 100 µm. (D) Morphology of N47 and PDFAT1 populations after 12 days of differentiation. Scale bar denotes 20 µm. Average BODIPY integrated intensity was multiplied by BODIPY count and divided by the number of nuclei. Statistical significance was assessed using two-way ANOVA with Tukey’s multiple comparison test. Hoescht shows genetic material and BODIPY shows intracellular neutral lipids. (*, **, ***, **** denote *p*□<□0.05, *p*□<□0.01, *p*□<□0.001 and *p*□<□0.0001, n=3).

### Consumers perceived distinct aroma differences between undifferentiated and differentiated cultivated fat

Sixty-one consumers were recruited from the Tufts University School of Arts and Sciences and School of Engineering student and staff populations. Consumers were given undifferentiated and day 12 differentiated porcine adipose tissue baked at 120°C for 45 mins in a commercial oven (Figure 2A). Adipocytes were differentiated until day 12 to increase biomass yield.

**Figure 2.**
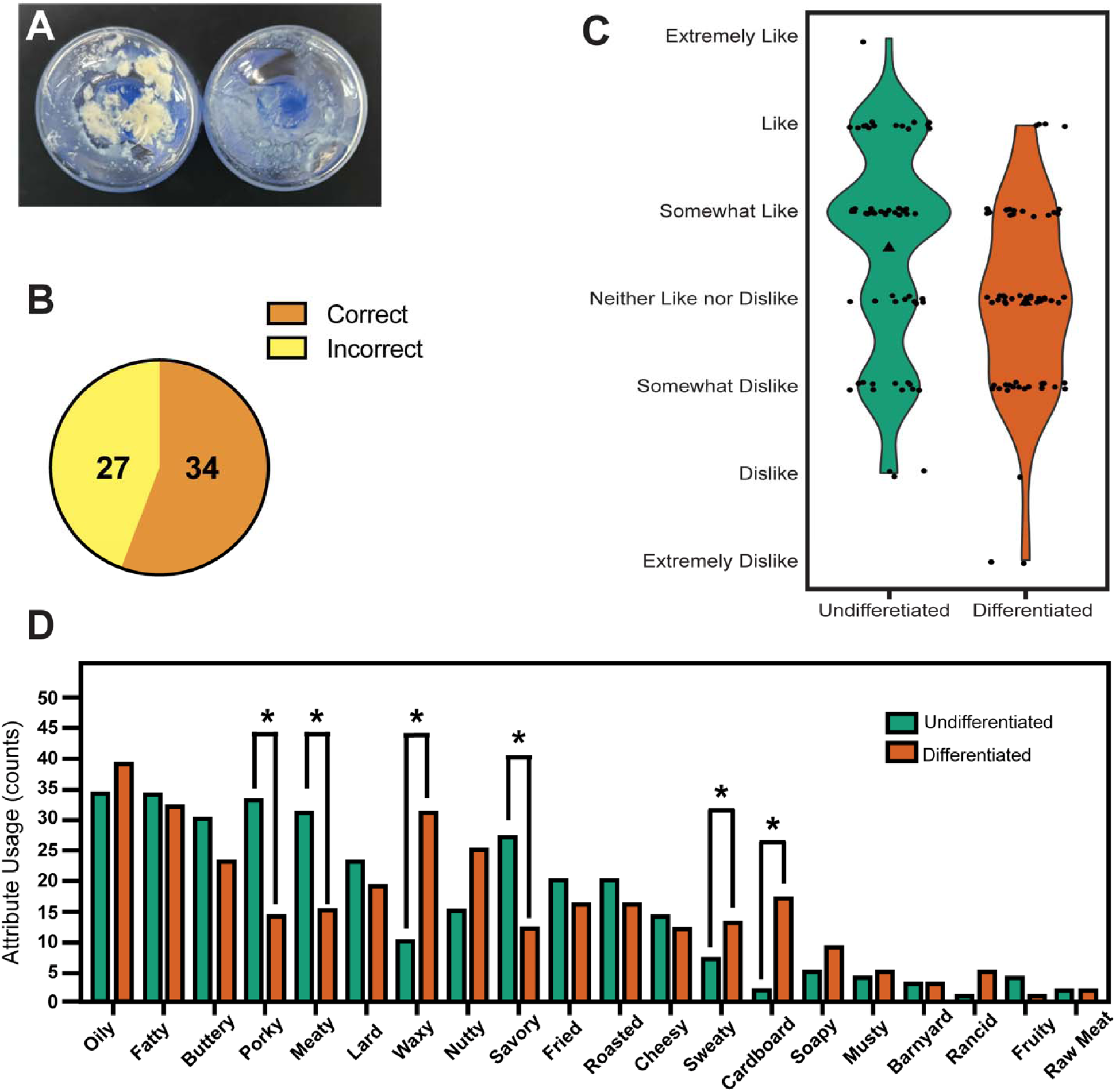
Results of sensory evaluation comparing differentiated and undifferentiated porcine adipose tissue. (A) image showing differentiated fat (left) and undifferentiated fat (right) after baking. (B) A panel of 61 consumer were able to discriminate between the two samples (C) Violin plot displaying the distribution of overall liking of aroma between differentiated and undifferentiated porcine fat samples based on 7-point hedonic scale of Extremely Dislike to Extremely Like. Mean hedonic rating is shown with a black triangle. (D) CATA results showing overall attribute usage by all participants. Sample groups were compared with Fischer’s exact test, where * and ** denote p□≤□0.05 and p□≤□0.01.

Consumers were presented with three fat samples in random combinations containing either two differentiated and one undifferentiated sample or one differentiated and two undifferentiated samples. In this triangle test, 34/61 consumers successfully discriminated between the samples (Figure 2B). Enough consumers were able to correctly discriminate to demonstrate statistical significance (p □<□0.05)^36^. Consumers also rated the overall aroma of the samples on a 7-point hedonic scale from “Extremely Dislike” to “Extremely Like” (Figure 2C). The violin plot displays hedonic ratings for each fat sample. Consumers expressed no difference in preference with respect to the aroma of differentiated and undifferentiated fat. When given a CATA task with aroma attributes “Oily”, “Fatty”, “Buttery”, “Porky”, “Meaty, “Lard”, “Waxy”, “Nutty”, “Savory”, “Fried”, “Roasted”, “Cheesy”, “Sweaty”, “Cardboard”, “Soapy”, “Musty”, “Barnyard”, “Rancid”, “Fruity”, and “Raw Meat”, consumers used attributes “Sweaty”, “Cardboard”, and “Waxy” with significantly higher frequency when smelling differentiated fat. Consumers used the attributes “Porky”, “Meaty” and “Savory” with higher frequency when smelling undifferentiated fat (Figure 2D).

### Adipogenesis leads to an increase in polyunsaturated fatty acids

N47 adipocytes were harvested at 0, 4, 8 and 12 days of adipogenesis via cell scraping. Lipids were extracted and converted to fatty acid methyl esters using transesterification for FAME analysis. Unsaturated fatty acids increased throughout differentiation and reached maximal detection at day 8 apart from arachidonic acid (20:4) (Figure 3A). Saturated fatty acids followed a similar pattern (Figure 3B). Fatty acids present in Intralipid supplementation were primarily linoleic acid, oleic acid, palmitic acid, alpha-linolenic acid, and stearic acid. Total fatty acid content also reached maximal detection at day 8 (Figure 3C). The most prominent fatty acids were 18:1n-9, 18:0, 16:0 (Figure 3D) and upon differentiation, 18:3n-3 was also detected at significant levels. Undifferentiated adipocytes contained a higher saturated to unsaturated fatty acid ratio with 56.9% saturated fatty acids and 43.1% unsaturated fatty acids. Starting at day 4 of adipogenesis, the saturated fatty acid proportion decreased to 26.5% from 56.9% in undifferentiated samples (Figure 3E).

**Figure 3.**
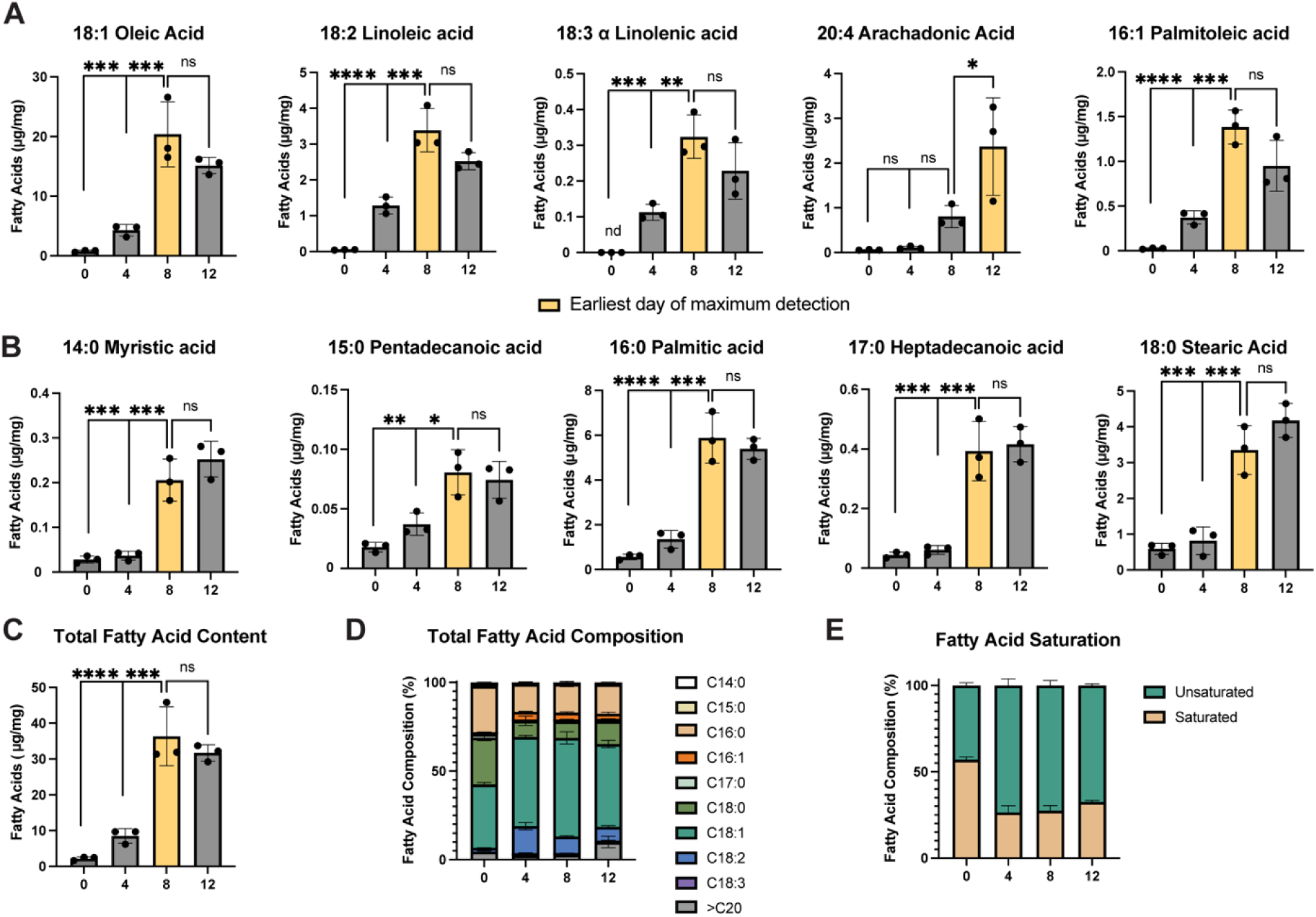
Analysis of fatty acid content throughout differentiation was conducted using FAME analysis with GC-FID. (A) Major unsaturated and (B) saturated fatty acids were detected in addition to (C) total fatty acid content and (D,E) total fatty acid composition. Yellow bars indicate the earliest day of maximum detection for a given fatty acid. Statistical significance was assessed using ordinary one-way ANOVA with Tukey’s multiple comparison test. (*, **, ***, **** denote *p*□<□0.05, *p*□<□0.01, *p*□<□0.001 and *p*□<□0.0001, n=3).

### Fatty aroma volatiles increase throughout differentiation

To assess differences in VOCs throughout differentiation, samples were analyzed with DHS-GC-MS. Due to maximized detection of fatty acid precursor molecules at day 8, fold changes of aroma volatiles were assessed between undifferentiated and day 8 adipocytes (Figure 4). The peaks of all compounds identified through deconvolution were normalized using the internal standard and cell mass after baking. Adipogenesis was associated with significant changes in the VOC profile of cultivated pork fat. Most enhanced lipid oxidation products between undifferentiated and day 8 adipocytes were furan containing molecules, ketones and aldehydes. Oxidation products with the largest fold changes were 2-pentylfuran (Figure 5A), 2-heptanone (Figure 5B) and aldehyde molecules hexanal and (E) 2-octenal (Figure 5C i, ii) which are known for their contributions to fatty, green and herbaceous aromas. Alongside 2-heptanone, methyl ketone molecules commonly found in dairy products such as 2-decanone and 2-nonanone and ketone derivatives such as maltol increased significantly. Other fatty aldehydes enhanced by differentiation include nonanal and pentanal. Significant enhancement was also observed in hydrocarbon molecules.

**Figure 4.**
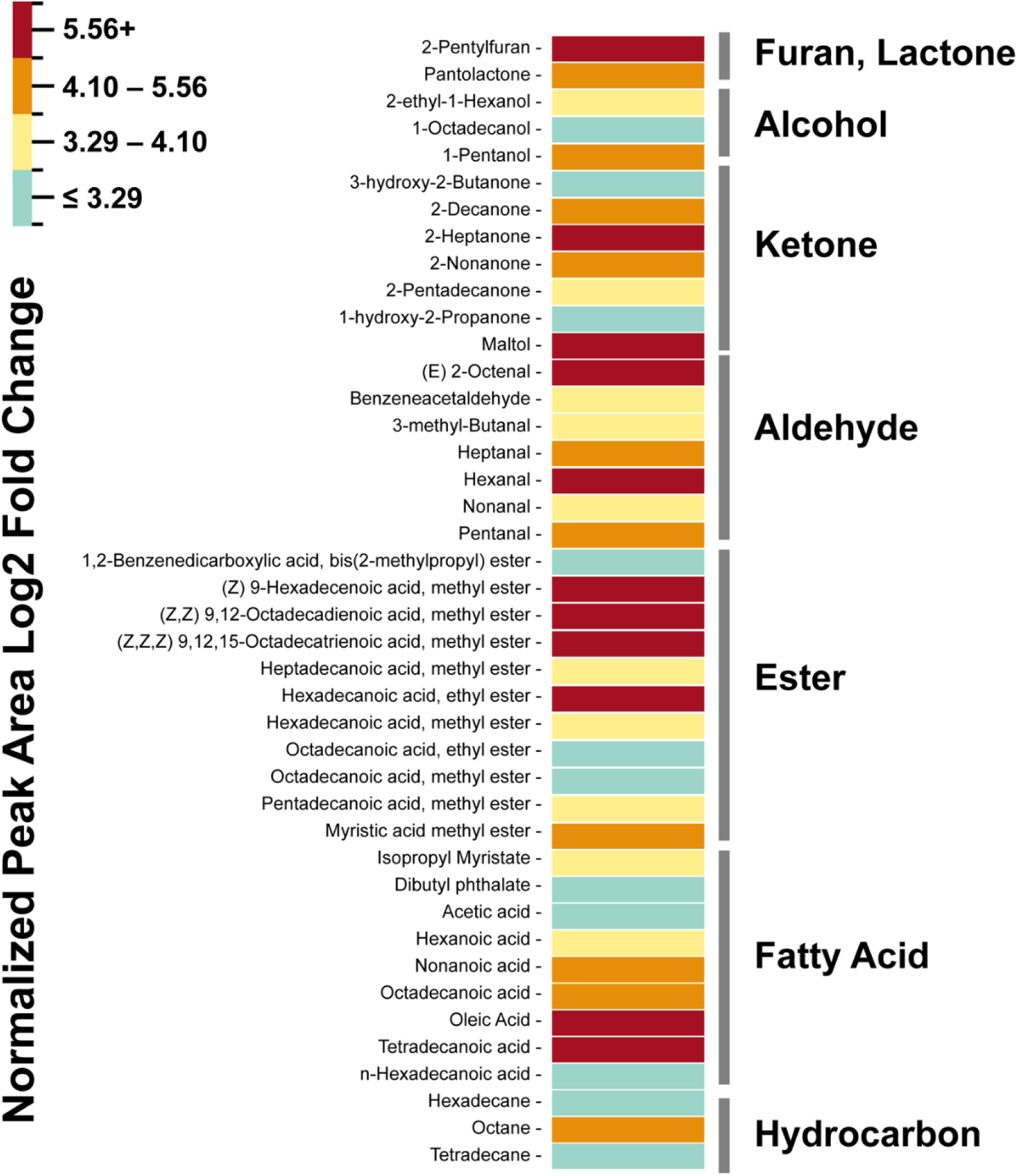
Total volatile profiling was conducted with DHS-GC-MS. The Log2 fold changes of the normalized peak areas of VOCs were calculated between undifferentiated and day 8 differentiated adipocytes. The color intensity represents the value of the Log2 fold changes grouped by quartiles where blue = bottom 25%, green = 25-50%, orange = 50-75% and red = top 25% of volatiles based on peak area enhancement, n=3. All VOCs are tentatively identified.

**Figure 5.**
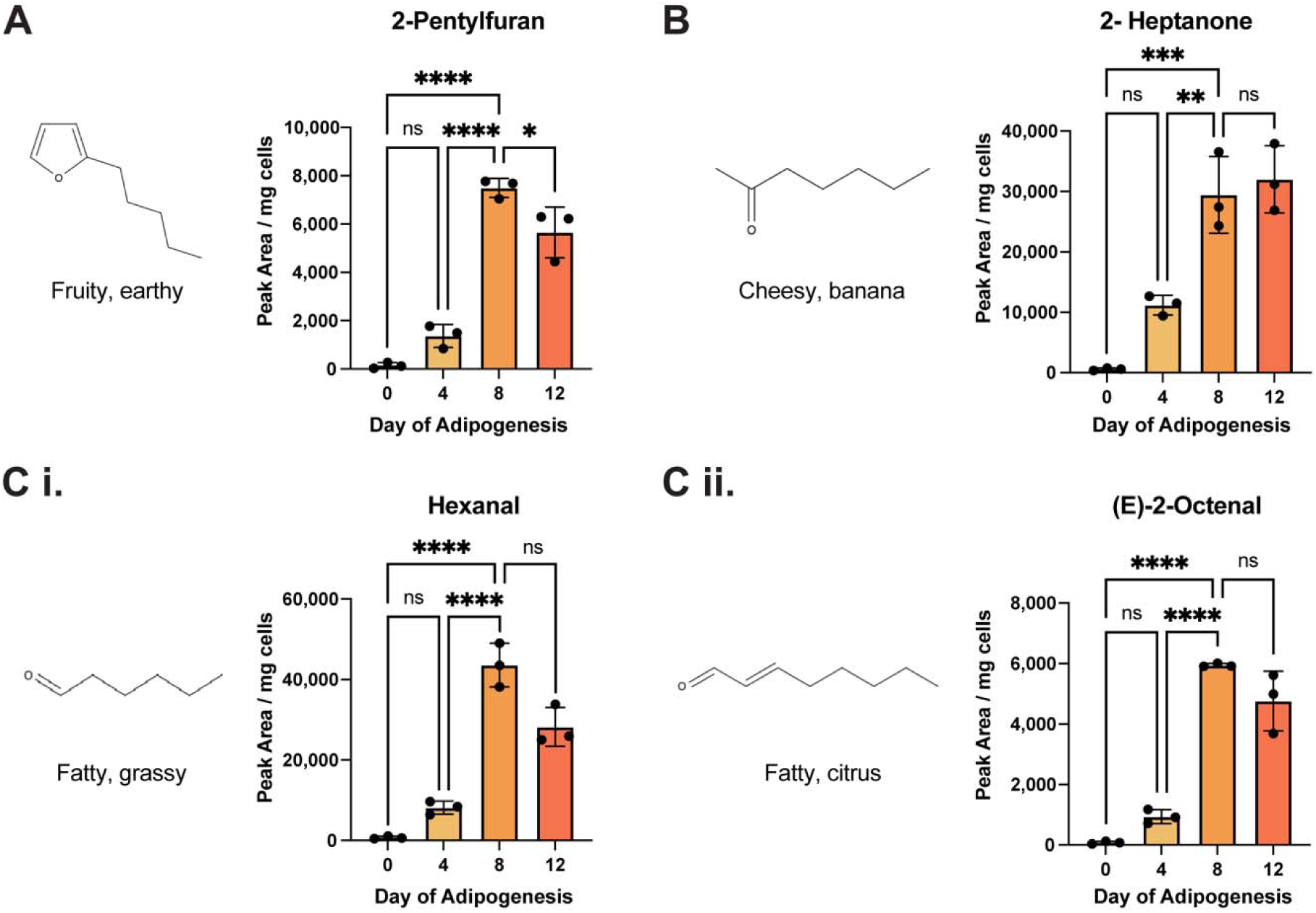
Volatile compounds with the largest fold changes outside of fatty acids and fatty acid methyl esters. (A) 2-pentylfuran (B) 2-heptanone and (C i,ii) aldehyde molecules hexanal and (E)-2-octanal were detected throughout adipogenesis. VOCs were quantified by normalizing peak areas to an internal standard and normalizing to cell mass after baking. Statistical significance was assessed using ordinary one-way ANOVA with Tukey’s multiple comparison test. (*, **, ***, **** denote *p*□<□0.05, *p*□<□0.01, *p*□<□0.001 and *p*□<□0.0001, n=3). Compounds were identified by retention index (RI) and Mass Spec referencing NIST17.

### Differentiated fat contributed key fatty aroma volatiles to conventional plant protein

To assess its contribution to key aroma volatiles in conventional plant proteins at low inclusion rates, cultivated fat was used as an additive to textured vegetable protein and VOC profiling was conducted with DHS-GC-MS. Defatted soy protein was blended with a traditional countertop blender and rehydrated with distilled water and 5% cultivated pork fat cells. Fat tissue was either

undifferentiated or differentiated for 8 days (Figure 6A). Again, lipid oxidation products showing the largest fold changes were furan containing molecules, ketones and aldehydes. With differentiated fat tissue as a supplement, 2-pentylfuran, 2-heptanone, hexanal and (E)-2-octenal were again among the most enhanced molecules (Figure 6B-D). Other molecules that were significantly increased included fatty aldehydes nonanal, hexadecanal and octadecanal, ketone molecules such as 3-hydroxy-2-butanone, and various fatty acid methyl esters and other ester molecules. No significant reduction of volatiles was observed for the addition of either DF or UD (Table 2). The TVP + DF samples showed a higher enhancement of fatty acid methyl esters and ethyl esters. The TVP + UD samples showed a lower enhancement of these molecules (Figure 6E).

**Figure 6.**
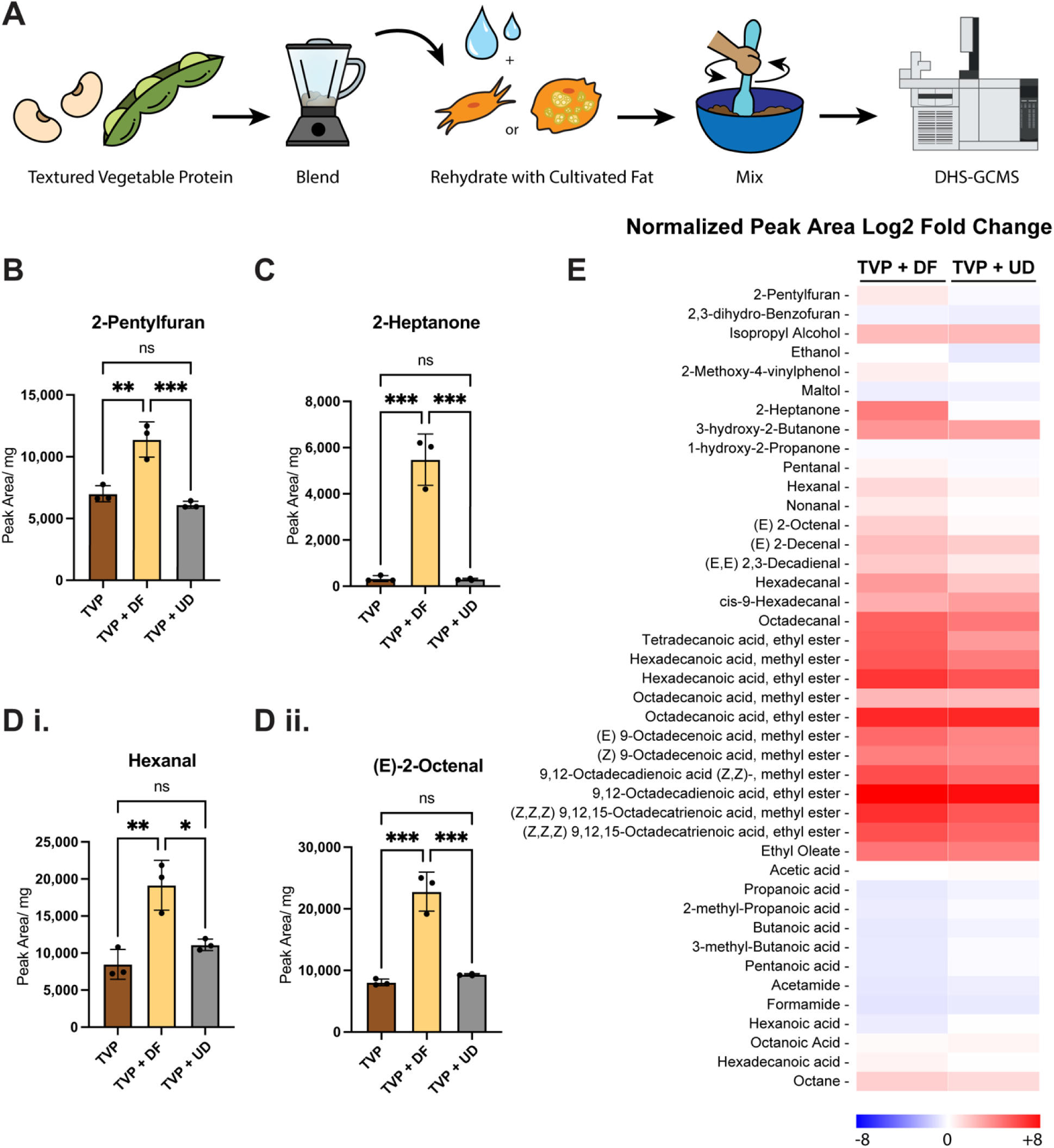
Assessment of volatile organic compounds from textured vegetable protein supplemented with day 8 differentiated (DF) and undifferentiated (UD) cultivated fat. (A) Samples were produced by blending commercially available TVP and rehydrating with water and 5% cultivated cells. Samples were mixed thoroughly before analyzing with DHS-GC-MS. (B) 2-pentylfuran (C) 2-heptanone and (D i,ii) aldehyde molecules hexanal and (E) 2-octanal were detected. (E) Log2 fold changes of normalized peak areas for compounds in TVP + DF or UD compared to TVP baseline. VOCs were quantified by normalizing peak areas to an internal standard and normalizing to cell mass after baking. Statistical significance was assessed using ordinary one-way ANOVA with Tukey’s multiple comparison test. (*, **, ***, **** denote *p*□<□0.05, *p*□<□0.01, *p*□<□0.001 and *p*□<□0.0001, n=3). Compounds were identified by retention index (RI) and Mass Spec referencing NIST17.

## Discussion

We demonstrated that fatty acid precursors and subsequent aroma volatiles in cultivated pork fat are altered significantly through adipocyte differentiation. Prior to assessing the volatile profile throughout adipogenesis, we compared the adipogenic capacity of our clonal cell line N47 and its parent population PDFAT1. We found that N47 achieved higher percent differentiation than PDFAT1 (Figure 2A,C) while both populations achieved their maximum percent differentiation at day 8. We also found on average, differentiated N47 cells were able to achieve more lipid accumulation than those in the PDFAT1 population (Figure 1B). This improvement was likely due to the homogeneity of the clonal population and previous selection for higher differentiating clones^37^. Similar to other clonal preadipocyte populations, N47 did not achieve 100% differentiation efficiency^38,39^, suggesting that further media optimization can be conducted (Figure 1C). Recent work in our laboratory identified the addition of basic fibroblast grown factor (bFGF) or a combination of A 83–01, CHIR99021 and Y-27632 (ACY) molecules in proliferation media significantly increased the growth rate and differentiation efficiency in PDFAT1^31^.

PDFAT was harvested in the proliferative state and after 12 days of differentiation to maximize biomass yield for the consumer sensory panel. Samples were cooked in a commercial oven at 120°C for 15 mins to align with DHS modules and standard cooking conditions. While this allowed for lipid oxidation and the release of VOCs, caramelization did not occur at this temperature (Figure 2A). A significant majority (34/61) of consumers determined that the aroma of differentiated PDFAT was unique to that of undifferentiated PDFAT (Figure 2B). This ratio was substantially lower than that observed in our previous comparison of cultivated and conventional pork adipose tissue^28^. This is likely due to the higher similarity of these samples in terms of production methods, as both were cultivated under comparable conditions using the same proliferation media and lacked competing cell types present in native tissue^40,41^. Although adipogenesis resulted in morphological changes that may influence aroma profiles, such as changes in extracellular matrix composition, collagen remodeling and decreased pericellular fibronectin as seen in 3T3-L1 adipocytes^42^, lipid accumulation was considered the most obvious factor associated with the observed differences in aroma^43–45^. Both trained and untrained panelists reliably identified the presence, absence, and in some cases, the type of lipids present in food samples^46–48^. While consumers did not report a significant preference for the aroma of one sample over the other, there was a trend towards “Somewhat Like” for undifferentiated cells and “Neither Like nor Dislike” for the differentiated cells, likely due to consumer preference for unoxidized fats when given oil-supplemented products (Figure 2C) ^49,50^. Consumers often associate “Cardboard”, “Waxy” and “Sweaty” attributes with aged fat or rancid oil aromas (Figure 2D)^51–53^. “Waxy” has been cited particularly for soybean oil, which is the primary component of Intralipid^54^. While consumers detected many attributes for warmed oil or lipids in differentiated fat, we hypothesize that due to lack of a muscle component such as free-amino acids and peptides, and sugar-degradation products formed during caramelization, particularly reactive carbonyl compounds, limited Maillard reaction products were scarcely volatilized and thus aroma attributes such as “Meaty” or “Porky” did not manifest to consumers. Additionally, previous studies have shown that higher lipid content can promote the retention of VOCs within the matrix. Accordingly, the diminished perception of “Meaty” and “Savory” aroma attributes in differentiated cells may be associated with increased lipid accumulation, which could have reduced the release of these compounds at this temperature^55^. Future sensory testing should be completed at higher temperature ranges to increase volatile formation and align consumer sensory perception with instrumental volatile profiles.

To investigate this difference in lipid content and fatty acid composition, fatty acid analysis was conducted throughout adipogenesis using GC-FID. Medium and long chain fatty acids serve as the primary precursor molecules to VOCs, particularly polyunsaturated fatty acids due to their susceptibility to oxidation by free radical and reactive oxygen species^56,57^. Despite observing an increase in differentiation efficiency and lipid accumulation through day 12, we found that all detected unsaturated fatty acids reached maximum detection at day 8, except for arachidonic acid which increased through day 12 (Figure 3A). This may occur due to large shifts in lipid metabolism in early-mid differentiation, where expression of oxidases and lipogenic enzymes are at their peak, and much smaller changes in total lipid content in late differentiation between day 8 and day 12^58,59^. Given that most fatty acid precursors appeared to reach their maximum levels by day 8, we hypothesize that day 8 adipocytes may provide flavor contributions broadly comparable to those of day 12 adipocytes. The main exception may be arachidonic acid, which can generate low-odor-threshold VOCs associated with egg-white-like and animal-like aroma notes^60^. These findings suggest that extending adipocyte culture beyond day 8 may offer limited additional flavor benefits, although further sensory and volatile analyses are needed to confirm this possibility. The same pattern was observed for saturated fatty acids (Figure 3B, 3C). Apart from palmitic acid and stearic acid which are present in Intralipid, we observed lower concentrations of saturated fatty acids than unsaturated. We also observed significant changes in overall fatty acid composition early in adipogenesis (Figure 3D, 3E). The highest contributing fatty acids were 18:1n-1, 16:0 and 18:0, which is in alignment with the composition of native pork fat which consists mainly of 18:1n-9 and 16:0^61,62^. Throughout differentiation, increases in ratios of 16:1, 18:2n-6 and 18:3n-3 fatty acids were observed. While undifferentiated cells primarily contained saturated fatty acids, by day 4 of differentiation the fatty acid profile shifted toward predominantly unsaturated species, likely driven by increased desaturase activity to support lipid storage and the incorporation of supplemented lipids^63,64^. These observations suggest that adipogenesis provides an opportunity to enhance flavor precursor formation while also modulating nutritional profiles for cultivated meat, particularly through increased monounsaturated and polyunsaturated fatty acids.

We examined whether adipogenesis would enhance the formation of lipid-derived aroma compounds commonly found in conventional meat. Adipogenesis significantly enhanced the formation of furans, lactones, alcohols, aldehydes, and select ketones (Figure 4). Notably, 2-pentylfuran which is a common product of linoleic acid decomposition and contributes to fruity and earth aromas in cooked fat, demonstrated a 50-fold increase in normalized peak area from undifferentiated cells^31,65^(Figure 5A). Additionally, ketones such as 2-heptanone which adds to floral and cheese aroma in cooked meat was particularly enhanced, with a 53-fold increase. The molecule 2-heptanone has also been found as a oleic acid degradation product that may contribute largely to the species-specific flavors of pork^23^ (Figure 5B). Fatty aldehydes such as hexanal and (E)-2-octenal where also among the most enhanced compounds (Figure 5C).

These compounds contribute to vegetative/grassy and citrus aromas to fat and showed a 59-fold and 76-fold increase, respectively^66–69^ (Table 1). Both hexanal and (E) 2-octenal are known linoleic acid and arachidonic acid degradation products^65,70,71^. Hexanal is associated with grassy and fresh aroma attributes at low concentrations but is often considered an off-odor associated with lipid rancidity in meat products^72,73^. In our previous study of conventional and cultivated pork fat tissue, the highlighted compounds were detected in both sample types. These volatiles were present with significantly higher peak areas in conventional pork fat compared to cultivated pork fat, with the exception of 2-heptanone which was present only in the cultivated samples^28^. Taken together, these findings suggest that differentiation is a key step toward matching cultivated pork’s volatile profile to that of conventional pork.

**Table 1.**
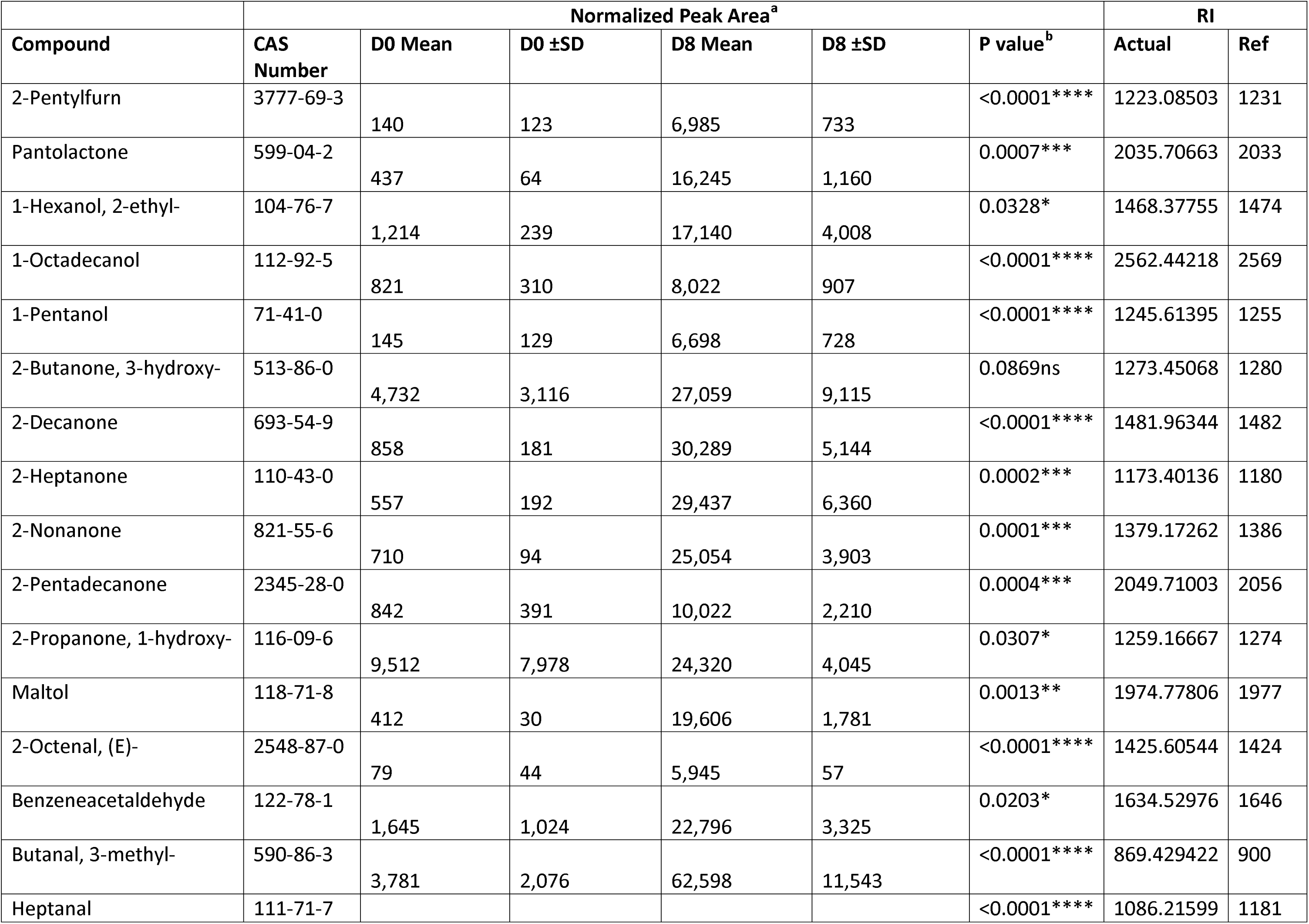
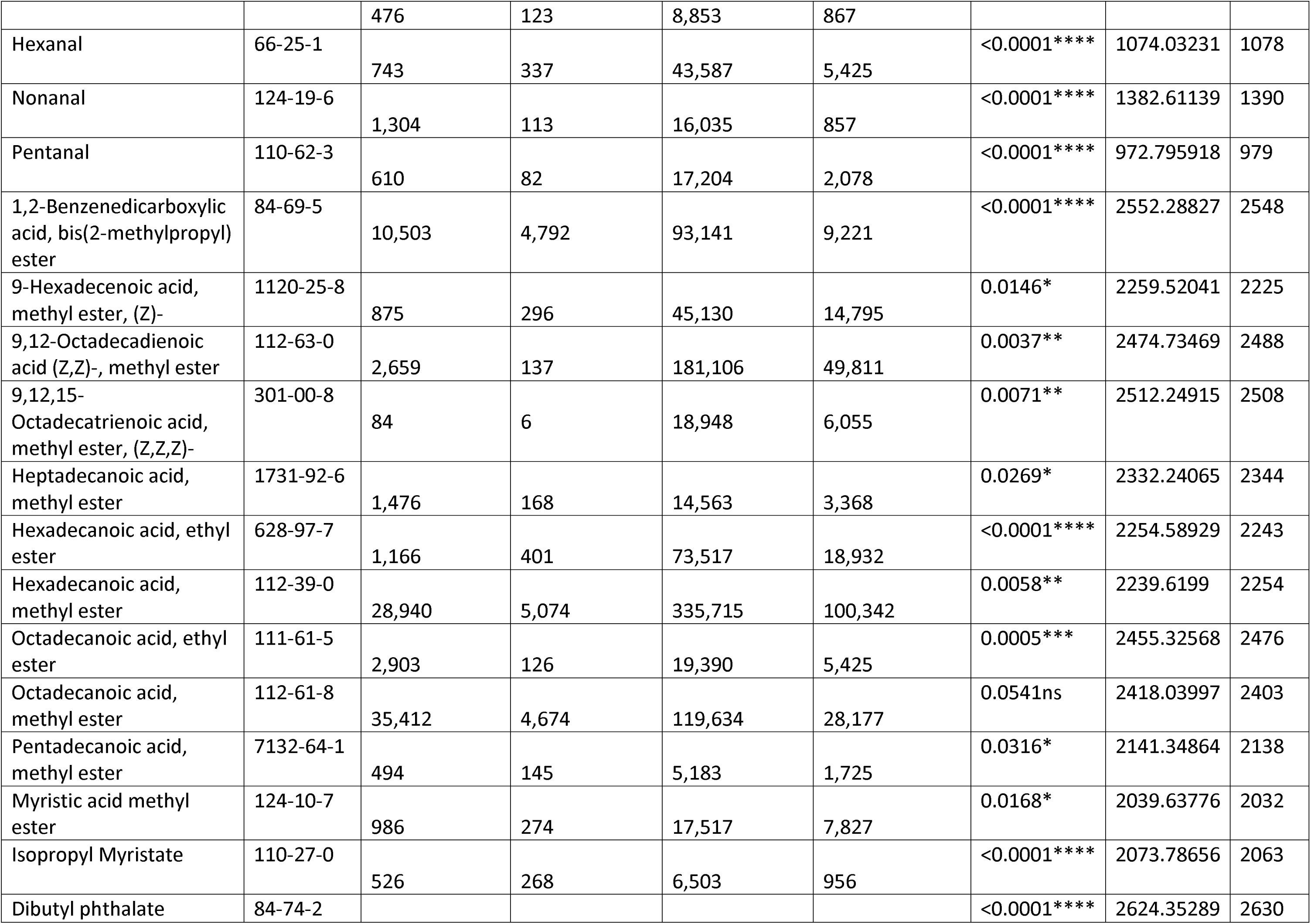
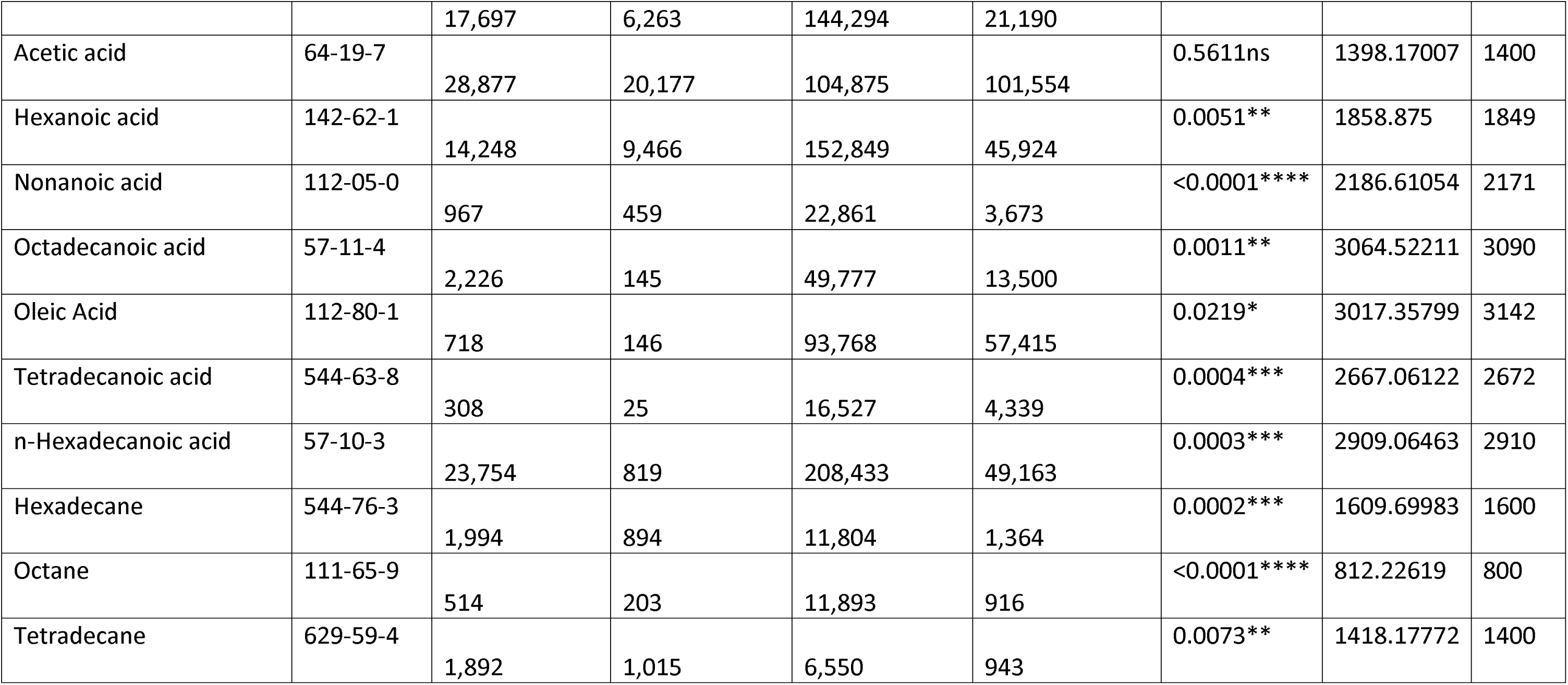
Volatile organic compounds (VOCs) detected from undifferentiated (D0) and differentiated (D8) cultivated porcine fat sampled at 120°C. (a) Average peak areas of compounds was normalized by internal standard Naphalene-D8 and by the dried cell mass (mg) after baking. (b) Statistical significance was determined using ordinary one-way ANOVA ( ns, *, **, ***, **** denote not significant, p⍰<⍰0.05, p⍰<⍰0.01 and p⍰<⍰0.001, p⍰< 0.0001 respectively). All VOCs were tentatively identified with mass spectrometry (MS) data and Retention Index (RI). RI references were identified in the literature from DB-Wax or standard polar column.

**Table 2.**
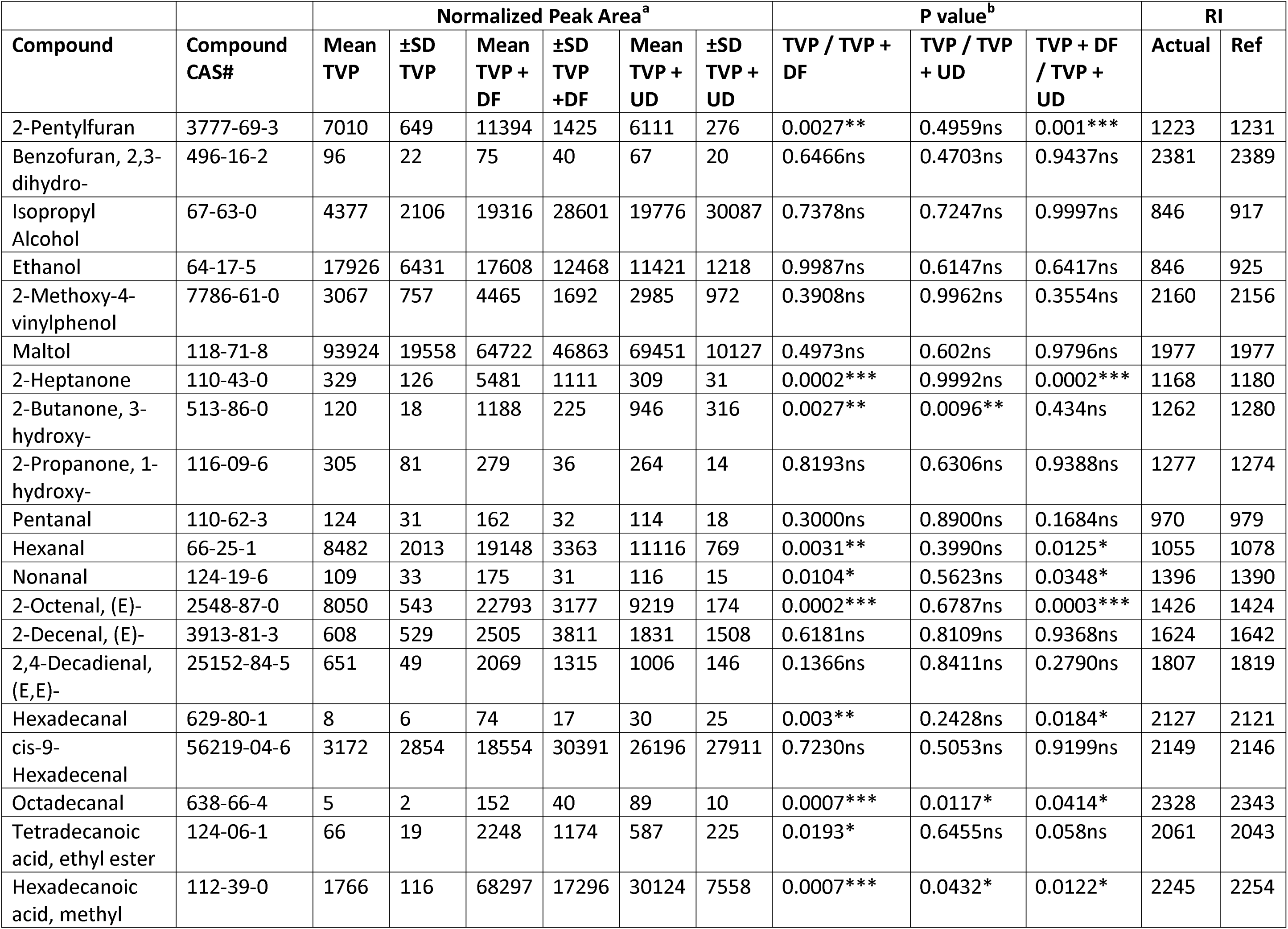
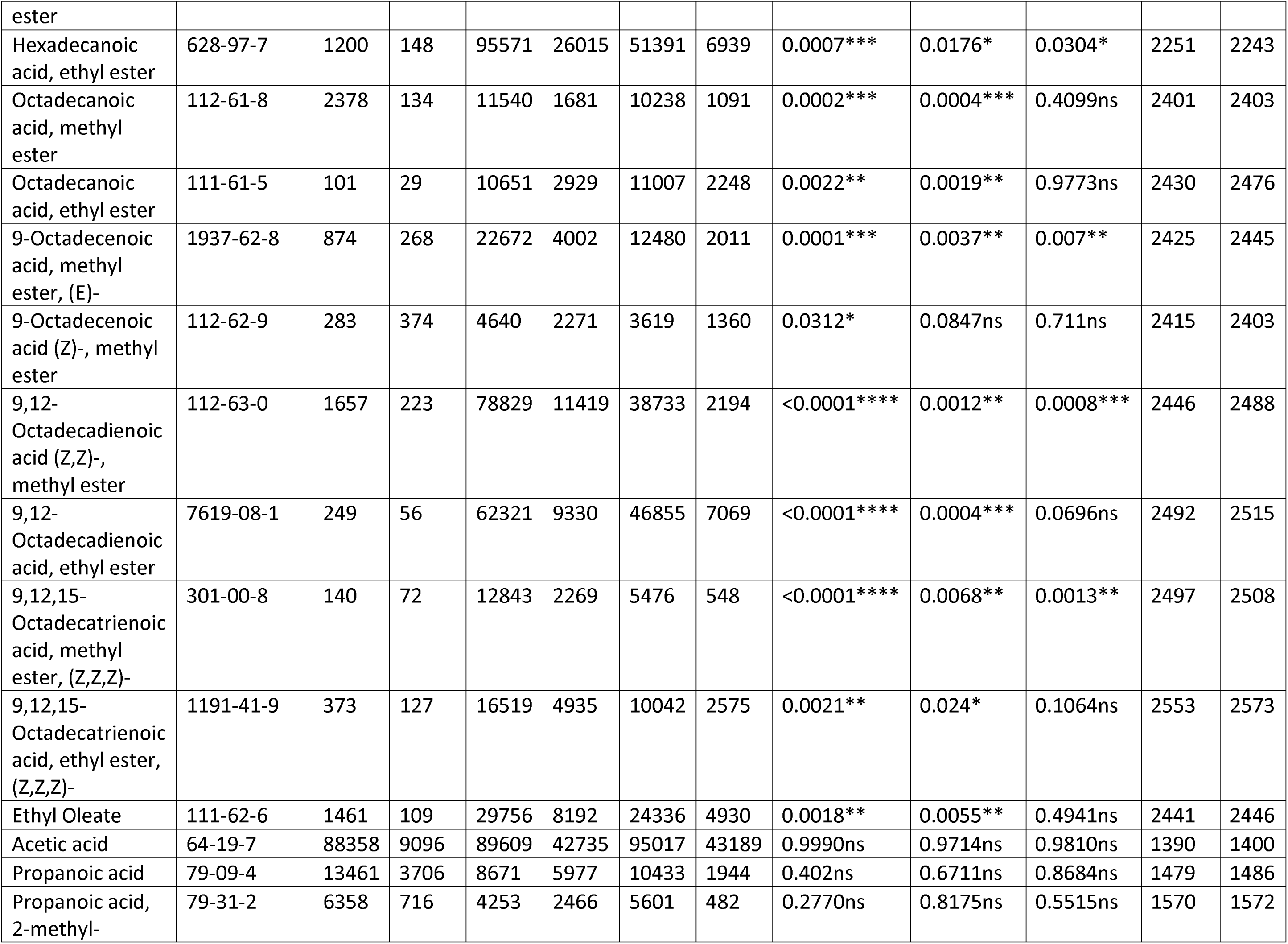
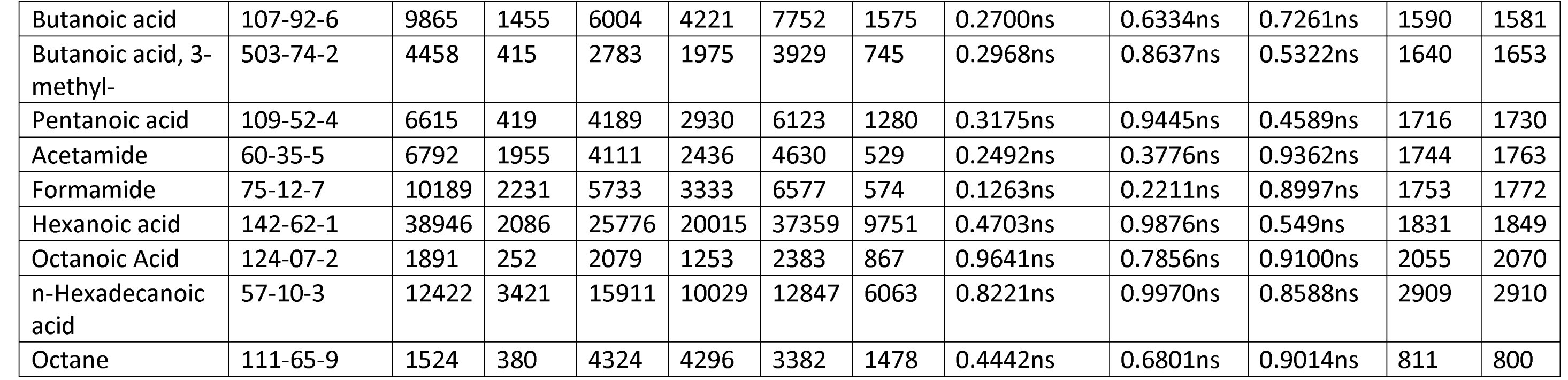
Volatile organic compounds (VOCs) detected from textured vegetable protein combined with undifferentiated (UD) and day 8 differentiated (DF) cultivated porcine fat sampled at 120°C. (a) Average peak areas of compounds was normalized by internal standard Naphalene-D8 and by the dried sample mass (mg) after baking. (b) Statistical significance was determined using ordinary one-way ANOVA ( ns, *, **, ***, **** denote not significant, p⍰<⍰0.05, p⍰<⍰0.01 and p⍰<⍰0.001, p⍰< 0.0001 respectively). All VOCs were tentatively identified with mass spectrometry (MS) data and Retention Index (RI). RI references were identified in the literature from DB-Wax or standard polar column.

Other largely enhanced ketone and aldehydes molecules included 2-decanone, 2-nonanone, 2-pentadecanone and pentanal which add sweet and green notes to fat^74^. Many aroma compounds associated with fattiness were enhanced, but our consumers found no difference in the “Fatty” attribute between the two samples. This may be because fattiness is an attribute that is strongly perceived through mouthfeel and taste in addition to odor, while attributes such as oiliness, which was detected by consumers in differentiated fat, is often reliably detected by odor alone^75,76^. Additionally, many fat-derived aldehydes possess low odor thresholds, potentially minimizing perceptible differences between the fat samples despite variations in their concentrations^77^. Hydrocarbon molecules such as hexadecane and tetradecane showed significant increases, but with smaller fold changes compared to ketones and aldehydes (Table 1).

To investigate the potential of cultivated fat as a flavor enhancer for plant-based proteins at low inclusion rates, defatted soy protein, a widely used textured vegetable protein, was combined with 5% (w/w) differentiated and undifferentiated adipocytes (Figure 6A). An inclusion level of 5% cultivated cells was selected to reflect commercially relevant hybrid formulations, where formulation details are often proprietary. Commercial precedent supports low inclusion levels, but these ranges are from 5–80% in regulatory submissions^78,79^. We analyzed the formation of lipid-derived aroma compounds as a result of the adipocyte inclusion and observed increases in furan, alcohol, ketone and fatty aldehyde molecules. Again, 2-pentylfuran, 2-heptanone, hexanal and (E)-2-octenal were significantly increased with the addition of differentiated adipocytes (Figure 6B-D). We observed a smaller 1.6-fold increase in 2-pentylfuran compared to plain TVP. Ketone molecule 2-heptanone was increased 3.4-fold and aldehydes hexanal and (E)-2-octanal increased 2.2-fold and 2.8-fold respectively. We also observed larger increases in aldehydes such as hexadecenal with an 8.8-fold increase (Table 2, Figure 6E). These molecules both contribute significantly to fatty flavor^80^. The addition of undifferentiated cells did not result in increases in concentration for these compounds. However, we observed select aroma molecules such as 3-hydroxy-butanone and some fatty acids whose concentrations were enhanced by undifferentiated cells alone (Table 2). This suggests that undifferentiated cells still provide some key flavor compounds, evident by positive sensory ratings to cultivated meat products made with fibroblasts or undifferentiated cell types^81^.

Differentiating cultivated adipocytes to produce lipid-laden fat serves as an approach for generating key aroma volatiles and providing nutritionally enhanced flavor supplementation for cultivated meat and alternative protein products. Fatty acid precursors were tunable during this process, enabling modulation of the resulting volatile profile which can be used to achieve species-specific or otherwise more desirable flavor profiles. This process can also be used to introduce media supplementation such as amino acids and vitamins to increase specific umami and deep fat aromas^31,82^. While differentiation requires additional media usage, cost, and time in culture, we find that maximal differentiation is not required to achieve the highest concentrations of aroma volatiles. This can be used advantageously to decrease adipocyte production times, and thus save significant costs for production. While differentiated adipocytes hold higher potential for flavor modulation of plant proteins, undifferentiated cells contributed select flavor compounds demonstrating that cellular components can contribute meaningfully to flavor generation. Thus, it is unclear whether the sole addition of exogenous oils or lipids to plant-based products is sufficient to fully replicate these flavor attributes. The present study finds adipocyte differentiation as a key step to producing cultivated meat or realistic meat alternatives.

## Methods

### Preadipocyte Cell Culture and Adipogenic Differentiation

Porcine DFAT cells were isolated from the belly (subcutaneous fat) of a 93-day-old female Yorkshire pig (DOB: 10/18/2021). Single cell isolation from porcine DFAT cells was conducted at the Tufts University School of Medicine Flow Cytometry Core by fluorescence-activated cell sorting (FACS) with the FACSAria (BD Biosciences, Franklin Lakes, NJ, USA)^28^. PDFAT1 and clone N47 were grown using high glucose Dulbecco’s modified eagle medium (DMEM) with GlutaMAX, phenol red, and sodium pyruvate (10569044; Thermo Fisher, Waltham, MA, USA)L+L20% Fetal Bovine Serum (FBS, A31606-01; Thermo Fischer)L+ 100 µg/ml Primocin (ant-pm-1; InvivoGen, San Diego, CA, USA) + 0.25 μg/cm^2^ laminin 511-E8 (N-892021; Iwai North America Inc., San Carlos, California, USA) in a 39°C, 5% CO_2_ incubator. General cell counting was performed using an automated cell counter (NucleoCounter NC-200; Chemometec, Lillerød, Denmark) and cell detachment was achieved enzymatically using TrypLE Express (12604013; Thermo Fischer). The cells were cryopreserved in 90% culture medium and 10% dimethyl sulfoxide (DMSO, D2438; Sigma).

Porcine preadipocytes were grown until confluency in proliferation media. After remaining confluent for at least 24 hours, cells were switched to adipogenic induction media consisting of DMEM + GlutaMAX with 10% FBS, 100 µg/ml Primocin, 10 µM biotin (B04631G; TCI America, Portland, OR, USA), 5.67 µM calcium-D-pantothenate (P001225G; TCI America), 3 µg/mL insulin (I0516; Sigma), 0.3 μM dexamethasone (DEX, AC23030; Thermo Fisher), 0.1 mM 3-isobutyl-1-methylxanthine (IBMX, I5879; Sigma), and 10 μM rosiglitazone (ROG, R0106; Thermo Fischer). Cells were then grown in lipid accumulation media containing DMEM + GlutaMAX, 3 µg/mL insulin, 10 µM biotin, 113 µM ascorbic acid (57-785-0; Fischer Scientific), and 500 µg/ml Intralipid (I141; Sigma). Cells remained in the lipid accumulation media for up to 10 additional days.

### Fat Harvest

Spent culture media was aspirated and adipocytes were rinsed with Dulbecco’s phosphate buffered saline with calcium and magnesium (DPBS, 1404013; Thermo Fischer) 3-5 times and flasks were kept vertical for 1-2 mins to thoroughly drain DPBS and remaining media. Once excess DPBS was aspirated, the adipocytes were harvested using a cell scraper (83.3952; Sarstedt, Nümbrecht, Germany). Periodically, additional liquid was gently aspirated from the flask. The cell mass was pushed to the back of the flask and collected, then stored at -80°C.

### Lipid Staining

Cultivated PDFAT1 and N47 adipocytes were stained to confirm intracellular lipid accumulation. Cells were rinsed with DPBS and fixed with 4% paraformaldehyde (PFA) for 20 min at room temperature. Cells were stained with DPBS supplemented with 2 µM BODIPY 493/503 (4,4-difluoro-4-bora-3a,4a-diaza-s-indacene, D3922; Invitrogen, Waltham, MA, USA) for intracellular lipids and 2 µg/ml Hoescht 33342 (H3570; Invitrogen) for nuclei. Cells were stained for 15-20 min at room temperature protected from light. Samples were rinsed with DPBS and imaged.

Imaging was performed with a fluorescent microscope (KEYENCE, BZ-X700, Osaka, Japan). Percent differentiation and differentiation efficiency was quantified by measuring normalized BODIPY intensity by the Celigo Image Cytometer (200-BFFL-5C; Nexcelom Bioscience LLC, Lawrence, MA). The differentiation percentage was calculated by BODIPY count divided by nuclei count. The average integrated intensity of BODIPY was multiplied by BODIPY count, then normalized by dividing by the nuclei count.

### Textured Vegetable Protein Samples

Textured vegetable protein (TVP, Bob’s Red Mill; Milwaukie, OR, USA) was blended using a traditional countertop blender. Rehydration was achieved with Gibco™ UltraPure distilled water (10977023; Thermo Fisher Scientific) at RT at a 1:2 ratio (w/w), with 5% cultivated adipocytes incorporated into the water before rehydration. Samples were mixed thoroughly to ensure uniform texture and left to rehydrate for 10-15 min. Samples of 50 mg were placed into 20 mL headspace vials (Restek, Bellefonte, PA) for GC-MS analysis.

### Fatty Acid Analysis

Lipid extraction was performed using methyl tert-butyl ether (MTBE) as described in our previous study^19^. Nonadecanoic acid (20 µL; 6 mg/mL in hexane; N5252, MilliporeSigma) was used as an internal standard. The MTBE phase was extracted twice, combined, and dried under nitrogen. The dried extract was saponified with 3 mL of 0.5 M sodium methoxide in methanol (92446; MilliporeSigma) at 55°C for 30 min, followed by methylation with 3 mL of 14% boron trifluoride in methanol (15716; MilliporeSigma) under identical conditions. At room temperature, the reaction mixture was transferred to a 15 mL centrifuge tube and mixed with 2 mL of saturated NaCl solution and 2 mL of hexane (139386; MilliporeSigma). Samples were vortexed and centrifuged at 3,500 rpm for 5 min. The upper organic phase was collected into autosampler vials (26590; Restek) for gas chromatography analysis.

Fatty acid composition was analyzed using an Agilent 6890N gas chromatograph (Agilent Technologies, Santa Clara, CA) equipped with a flame ionization detector (FID) and a FAME capillary column (CP7430; 100 m × 0.25 mm i.d. × 0.25 µm film thickness; Agilent Technologies). Samples (1 µL) were injected at 250 °C with a split ratio of 1:20, using helium as the carrier gas. The oven temperature program consisted of an initial hold at 100°C for 5 min, followed by a ramp at 10°C/min to 220°C with a 28 min hold, and a ramp to 250°C with a 10 min hold. Fatty acids were identified by comparison of retention times with a commercial standard mixture (Food Industry FAME Mix; 35077; Restek, Bellefonte, PA).

### Dynamic Headspace GC-MS

Thermal desorption unit (TDU) tubes were packed with Tenax resin (11982; MilliporeSigma) and conditioned at 300 °C for 120 min under a constant flow of ultra-pure nitrogen. Following conditioning, 1 µL of naphthalene-d8 (31043; Restek), prepared at 10 µg/mL in dichloromethane, was injected directly into the Tenax resin. Residual dichloromethane was removed by reconditioning the tube at 75°C for 5 min under ultra-pure nitrogen. Naphthalene-d8 was used as an internal standard to normalize GC/MS injection efficiency. Cultivated fat samples were thawed from -80°C and weighed at RT. Next, 100 mg of wet sample was placed into 20 mL headspace vials, which were transferred to the dynamic headspace (DHS), equilibrated at 120°C, and incubated for 15 min. Following incubation, vials were purged with 200 mL of ultra-pure nitrogen at a flow rate of 100 mL/min. Preconditioned Tenax TDU tubes were then installed in the DHS trap module and maintained at 40 °C. Headspace sampling was performed by incubating the vial at 120°C and purging with 1,500 mL of ultra-pure nitrogen at 50 mL/min, with volatile compounds trapped on the Tenax® resin at 40°C. After trapping, a dry purge was conducted using 750 mL of ultra-pure nitrogen at 100 mL/min while maintaining the trap at 40°C. Following DHS collection, the TDU tube was transferred to the thermal desorption unit. Prior to desorption, the cooled injection system (CIS) equipped with a glass bead liner was cooled to −120°C using liquid nitrogen. The programmable temperature vaporizing (PTV) inlet was set to a split ratio of 1:10, initiating solvent purge at 0.01 min with a flow rate of 12 mL/min. Thermal desorption was achieved by ramping the TDU from 40°C to 250 °C at 720°C/min, followed by a 3 min hold at 250°C.

Chromatographic analysis was initiated by rapidly heating the CIS from −120°C to 250°C, followed by a 3 min hold. Separation was performed on an Agilent 7890A gas chromatograph equipped with a DB-WAX UI capillary column (30 m × 0.25 mm i.d., 0.25 µm film thickness; Agilent Technologies). The oven temperature program consisted of an initial hold at 40°C for 2 min, followed by a ramp of 5°C/min to 250°C and a final hold at 250°C for 15 min. Helium was used as the carrier gas at a constant flow rate of 1.2 mL/min. Mass spectrometric detection was performed using an Agilent MSD (−70 eV). The ion source and quadrupole temperatures were set to 250°C and 150°C, respectively. A solvent delay of 1.25 min was applied, and data were acquired in scan mode over a mass range of m/z 35–450.

### Sensory Panel Recruitment

Sixty-one consumers were recruited from the School of Arts and Sciences and the School of Engineering at Tufts University in Medford, Massachusetts. Recruitment was conducted through flyer advertisements and university email lists. Participants were informed of the nature of the study, as well as any potential risks or benefits associated with participation. Eligibility criteria included being at least 18 years of age and willingness to evaluate the aroma of cultivated meat products. Participants were assumed to be untrained, with no prior experience in triangle tests or check-all-that-apply (CATA) sensory evaluation methods.

### Triangle Test

Fat samples were stored at −80°C and thawed prior to analysis. Aliquots (100 mg) were transferred to headspace vials and heated at 120°C for 45 min in a commercial oven to generate aroma. Throughout the sensory evaluation, sample vials were maintained in bead baths at 60°C. The study followed guidelines outlined by the American Society for Testing and Materials (ASTM) in the Standard Guide for Serving Protocol for Sensory Evaluation of Foods and Beverages (E1871−17), which recommends a minimum holding temperature of 57 °C to minimize microbial growth. Preliminary testing was conducted to assess aroma stability at the 60°C holding temperature for up to 6 hours. No detectable changes in odor or development of rancid off-odors were observed during this period that would interfere with an untrained consumer’s ability to complete the discrimination task. Data were collected over a single day during a 6-hour period, consisting of five sessions with a maximum of twelve consumers per session. In total, 61 consumers participated. Consumers completed a triangle test designed to differentiate between the aroma of differentiated and undifferentiated fat. Each participant evaluated one triangle set consisting of three samples, presented in randomized order. All six possible sample combinations (AAB, ABA, BAA, BBA, BAB, and ABB) were distributed across the study. Sample vials were concealed to mask visual differences between adipose tissues.

### Consumer Preference and CATA

Sample preparation for the check-all-that-apply (CATA) analysis was identical to that used for the triangle test. The same 61 consumers participated in the CATA analysis and provided preference data using a questionnaire that included a 7-point hedonic scale in combination with a CATA task. Two fat samples were presented in random order for every consumer. The ballot questions were as follows:

● Two questions using the 7-point hedonic scale ranging from Extremely Dislike to Extremely Like: “Please indicate your liking of the aroma found in Vial X.”
● Two check-all-that-apply (CATA) questions: “Which attributes would you use to describe the aroma found in Vial X? Check all that apply” from a list of the following descriptors: Oily, Fatty, Buttery, Porky, Meaty, Lard, Waxy, Nutty, Savory, Fried, Roasted, Cheesy, Sweaty, Cardboard, Soapy, Musty, Barnyard, Rancid, Fruity, Raw Meat.

### Statistical Analysis

Statistical analyses were conducted GraphPad Prism 10.6.1 and R version 4.5.2. Analyses include Analysis of variance (ANOVA) with Tukey’s post-hoc tests and Fisher’s exact tests. Error bars and ± ranges represent standard deviations. A p-value of 0.05 was used for statistical significance. All experiments in this study were carried out with at least triplicate technical samples (n≥3).

## Author Contributions

ETL: Conceptualization, Methodology, Investigation, Formal analysis, Data curation, Visualization, Writing – original draft and editing. NS: Methodology, Investigation, Visualization, Writing – review & editing. SM: Investigation, Writing: review & editing. SCF: Methodology, Formal Analysis, Visualization, Supervision, Writing – review & editing. DLK: Conceptularization, Supervision, Resources, Funding Acquisition, Writing – review & editing.

## Competing Interests

Author NS is employed by Ajinomoto Co., Inc. but declares no non-financial competing interests. All other authors declare no financial or non-financial competing interests.

## Acknowledgments

This study was funded by United States Department of Agriculture (2021-69012-35978). The funder played no role in study design, data collection, analysis and interpretation of data, or the writing of this manuscript. We thank the Tufts University Center for Cellular Agriculture (TUCCA). We acknowledge the Tufts Comparative Medicine services (CMS) for providing porcine adipose samples. We thank Anson Kwan for his assistance with formal analysis.

Figures were created using Adobe Illustrator 2026 version 30.0. Molecular structures were designed by MolView.

## Ethics Declaration

Isolation of adipocyte progenitor cells from a 93-day-old female Yorkshire pig (*Sus domesticus)* was approved by the Tufts University Medical Center and sourced from the Tufts Comparative Medicine Services. CMS methods are in accordance with the guidelines of the United States Department of Agriculture (USDA), Office of Laboratory Animal Welfare (OLAW), Massachusetts Department of Public Health (MDPH) and Association for Assessment and Accreditation of Laboratory Animal Care International (AAALAC). Additionally, all methods were conducted in accordance with ARRIVE guidelines 2.0.

The sensory evaluation study was approved by the Tufts University Health Sciences Institutional Review Board (IRB) for research involving human subjects (Study 00003031). All experiments were performed in accordance with the Tufts University Health Sciences IRB guidelines.

## Data Availability

All data generated or analyzed during this study are included in this published article. Data corresponding to human subjects research is not publicly available and not available upon request due to confidentiality of data under the Tufts University Health Sciences IRB.

